# The Topology of Mutated Driver Pathways

**DOI:** 10.1101/860676

**Authors:** Raouf Dridi, Hedayat Alghassi, Maen Obeidat, Sridhar Tayur

## Abstract

Much progress has been made, and continues to be made, towards identifying candidate mutated driver pathways in cancer. However, no systematic approach to understanding how candidate pathways relate to each other for a given cancer (such as Acute myeloid leukemia), and how one type of cancer may be similar or different from another with regard to their respective pathways (Acute myeloid leukemia vs. Glioblastoma multiforme for instance), has emerged thus far. Our work attempts to contribute to the understanding of *space of pathways* through a novel topological framework. We illustrate our approach, using mutation data (obtained from TCGA) of two types of tumors: Acute myeloid leukemia (AML) and Glioblastoma multiforme (GBM). We find that the space of pathways for AML is homotopy equivalent to a sphere, while that of GBM is equivalent to a genus-2 surface. We hope to trigger new types of questions (i.e., allow for novel kinds of hypotheses) towards a more comprehensive grasp of cancer.

## 1 Introduction

Let us begin by recalling a quote of Henri Poincare:

> Science is built up of facts, as a house is built of stones; but an accumulation of facts is no more a science than a heap of stones is a house.

Cancer is driven by somatic mutations that target signaling and regulatory pathways that control cellular proliferation and cell death [25]. Understanding how this happens is of paramount importance in order to improve our ability to intervene and attack cancer. Since the advent of DNA sequencing technologies, our understanding has progressed enormously and resulted in useful therapies.^1^

Notwithstanding the above, cancer morbidity is still very high and our understanding is still incomplete. Certainly, finding more relevant pathways will be helpful. Indeed we have developed novel formulations and algorithms that are computationally effective in doing so, a subject of our companion paper ([1] that builds on our work [2, 3]). We also believe that to start building a house it is not enough to simply accumulate more stones faster. To that end, in this paper, we look at whether the stones that we have identified so far form patterns that can help us build a house, using methods from algebraic topology (see Table 1).

**Table 1:** Correspondence between pathways and algebraic topology.

| Pathways | Algebraic topology |
| --- | --- |
| Regulatory/signaling pathway | simplex |
| The Emergent integrated circuit of the cell | simplicial complex $K_\varepsilon$ |
| Number of patients shared between two genes | persistent parameter $\varepsilon$ |

The view we are taking here is also articulated in *The Hallmarks of Cancer* ([7]):

> Two decades from now, having fully charted the wiring diagrams of every cellular signaling pathway, it will be possible to lay out the complete “integrated circuit of the cell” upon its current outline (Figure 1). We will then be able to apply the tools of mathematical modeling to explain how specific genetic lesions serve to reprogram this integrated circuit… One day, we imagine that cancer biology and treatment–at present, a patchwork quilt of cell biology, genetics, histopathology, biochemistry, immunology, and pharmacology–will become a science with a conceptual structure and logical coherence that rivals that of chemistry or physics.

**Figure 1:**
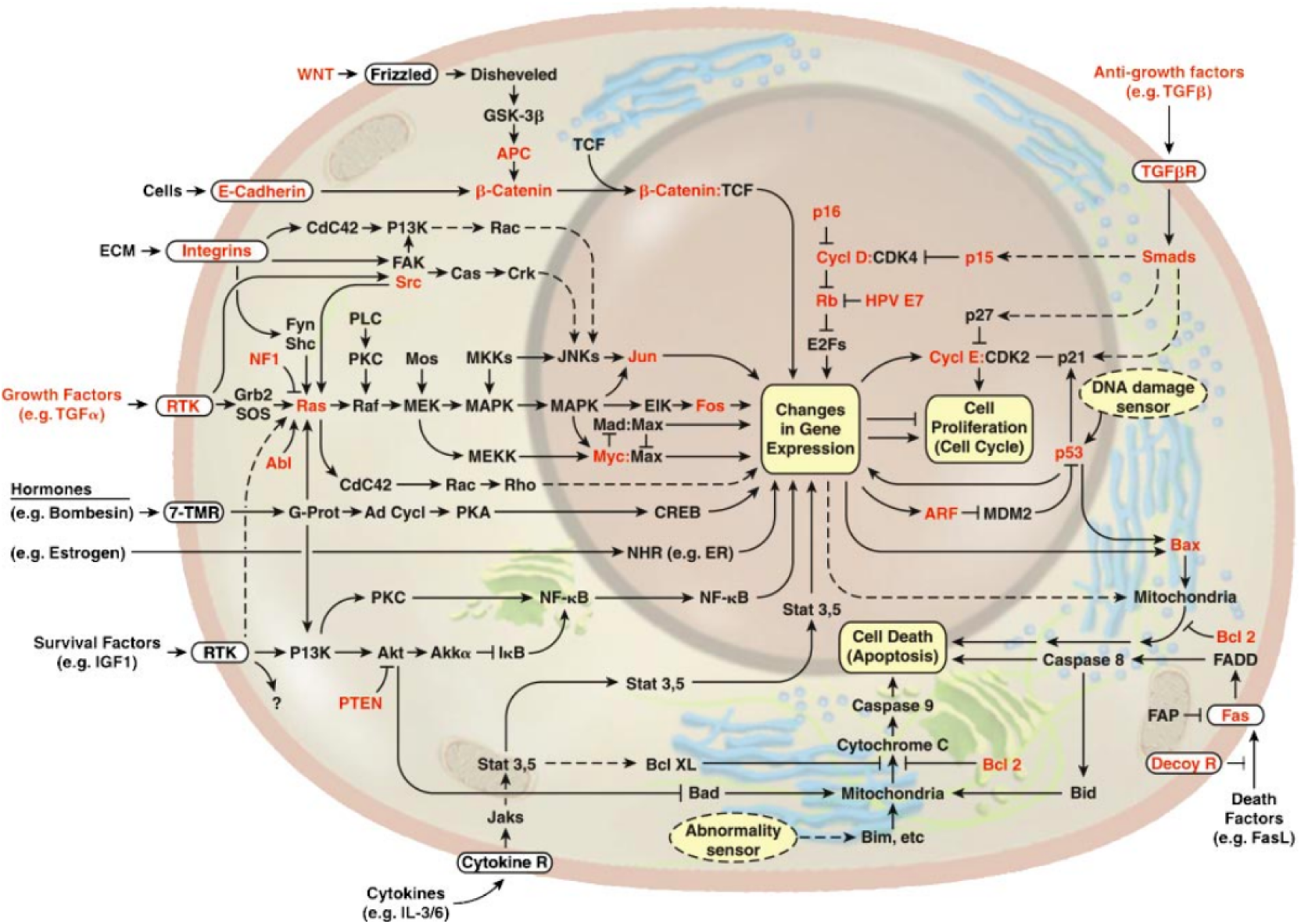
Cell circuitry, wiring pathways, resembles electronic integrated circuits. A full understanding of this picture will, one day, replace the present day patchwork with a more rational approach based on conceptual structures and logical coherence similar to those encountered in chemistry and physics [7].

Our topological journey departs from the (well trodden) path that recognized that signaling or regulatory pathways can be viewed as *independent sets* (modulo some notion of tolerance) in a matrix with rows as patients and columns as genes, which is used in determining new pathways via different computational methods (see our companion paper [1] and previous works [21, 22, 5, 18, 10]). Our key insight is that these pathways (however discovered), when grouped together, define a *simplicial complex* which is, pictorially, a polytope with faces given by those pathways. This vantage point connects us to the marvellous world of topology where simplicial complexes are the prototypes of spaces with shapes. Our journey explores this *notion of shape* in cancer genomics.

Our framework is built on this sequence of assignments:

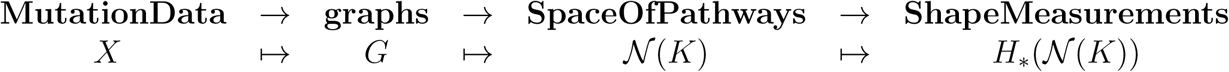

where:

- *G* is the mutation graph. That is, is the graph defined by the set of genes in *X* where two genes are connected if they have harboured mutations for the same patients. *K* is the simplicial complex given by the set of all independent sets of *G*. The underlying mathematics detailed in [6] is based on the so-called Mayer-Vietoris construction, which itself is articulated around clique covering the graph *G*. We prove in [6] that, for a large class of graphs, such coverings provide most of the simplices of *K*.
- 𝒩(*K*) is the space of pathways. The facets (maximal independent sets) of *K* are the pathways, and their nerve defines the space of pathways 𝒩(*K*). Pictorially, the space of pathways is visualized through its 1-skeleton of the graph with pathways as vertices and where two pathways are connected by an edge if they intersect.
- *H*_*_(𝒩(*K*)) is the homology of the space of pathways, that is, the shape measurements of the space of pathways.

We have applied our approach to two different mutation data obtained from TCGA: Acute myeloid leukemia (AML) [14] and Glioblastoma multiforme (GBM) [13]. For both the data, we have computed the assignment

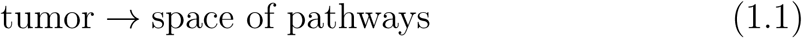

using persistent homology.^2^ Our calculation shows that the space of pathways for AML mutation data is homotopy equivalent to a sphere, while in the case of GBM data, the space of pathways is homotopy equivalent to figure eight (genus-2 surface).

## 2 Related works

The motivation for the present work originates from the work of the Raphael Lab, centred around the Dendrix algorithm [23], and its later improvements including CoMEt [9]. Both algorithms are in widespread use in whole-genome analysis–for instance, in [16, 17, 4, 20]. Building on those foundations, our work extends in the following two directions: First, the two key notions of exclusivity and coverage are abstracted here by the two simplicial complexes *K*_*ε*_ and *K*_*η*_ or, more precisely, by the filtrations of simplicial complexes 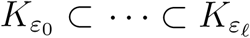 and 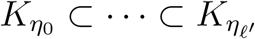, where we allow the two parameters to vary: *ε*_0_ ≤ *ε* ≤ *ε*_*ℓ*_ and *η*_0_ ≤ *η* ≤ *η*_*ℓ*′_. We do so because, in reality, the assumptions that driver pathways exhibit both high coverage and high exclusivity need not to be strictly satisfied. The functoriality of persistent homology, which takes the two filterations as input, handles this elegantly; we decide on the values of these two parameters after computing the barcodes. This type of analysis is not present in the aforementioned works. Second, motivated by the *naturality* of the constructions, the present paper goes beyond the computational aspects and ventures into the *conceptuality of cancer*. We have introduced the notion of the topological space of pathways 𝒩, together with its homology spaces, as a paradigm to rationalize the extraordinary complexity of cancer. To the best of our knowledge, this is a “provocative” idea that has not been explored before.

Another related set of works is [11] and [15] (from Stanford’s applied topology group), which also use algebraic topology tools in cancer. However, they differ from ours on two counts: the nature of the problem treated and the methodology used. The algorithm introduced, called Mapper, is a topological clustering algorithm, and it is not based on persistent homology. Mapper was used to identify a subgroup of breast cancers with excellent survival, solely based on topological properties of the data.^3^

## 3 Connecting to algebraic topology

Different errors occurring during data preparation (i.e., sequencing step, etc.) affect the robustness of the results. This implies that the computed pathways are likely to be affected by these errors and can not be considered as a robust finding without explicitly modeling the error in our constructions. To that end, the assignment we mentioned above is done as follows:

1. We think about the number of patients that are shared between two genes as a parameter *ε* (thus, absolute exclusivity corresponds to taking this parameter to zero).
2. Instead of applying our procedure once, we apply it for a range of values of the exclusivity parameter *ε*. That is, we consider a filtration of graphs (instead of one):

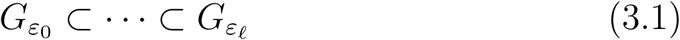

where for each graph *G*_*ε*_ two genes are connected if they have harboured mutations concurrently for at least *ε* patients. This yields a second filtration of simplicial complexes (we call such a filtration, a *persistent pathway complex*)

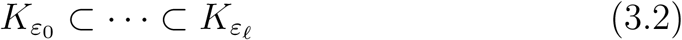

where *K*_*ε*_ is one of the three complexes we define below.
3. Measure the shape (the homology) of the different pathway spaces and then “average” the shape measurements that are obtained.

This (practical) version of homology is what we refer to as persistent homology. It tracks the persistent topological features through a range of values of the parameter; genuine topological properties persist through the change of the parameter whilst noisy observations do not (all of these will be made precise below). The key point is that the mapping *G*_*ε*_ ↦ *K*_*ε*_ is *functorial*; that is, it sends a whole filtration (i.e., 3.1) into another filtration (i.e., 3.2). In other words, it is not only sending graphs to simplicial complexes but it is also preserving their relations. This functoriality is at the heart of persistent homology.

### 3.1 The Gene-Patient graph

Consider a mutation data for *m* tumors (i.e., patients), where each of the *n* genes is tested for a somatic mutation in each patient. To this data we associate a mutation matrix *B* with *m* rows and *n* columns, where each row represents a patient and each column represents a gene. The entry *B*_*ig*_ in row *i* and column *g* is equal to 1 if patient *i* harbours a mutation in gene *g* and it is 0 otherwise. For a gene *g*, we define the fiber

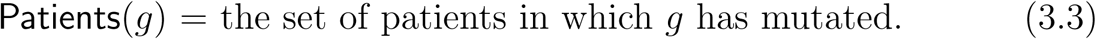

#### Definition 1

*The* mutation graph *associated to B and ε* > 0 *is the graph G*_*ε*_ *whose vertex set is the set of genes and whose edges are pairs of genes* (*g, g*′) *such that*

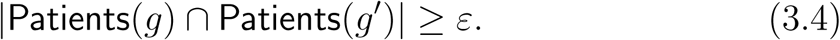

There are evidences that pathways can be treated as independent sets of mutation graphs (although not stated graph-theoretically) [24, 26].

### 3.2 The space of pathways

We would like to assign to the mutation graph an independence complex. We present below two different functorial ways to do so.

#### Definition 2

*Given a mutation graph G*_*ε*_, *its* persistent pathway complex *K*_*ε*_ *is the independence complex of G*_*ε*_ *(or equivalently, the clique complex of* 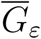, *the complement of G*_*ε*_).

We also define the persistent pathway complex *K*_*η*_.

#### Definition 3

*The persistent pathway complex K*_*η*_ *is defined as follows. Fix ε* = *ε*_0_ *and let* 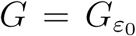. *The complex K*_*η*_ *is the complex generated by all independent sets S of G with coverage*

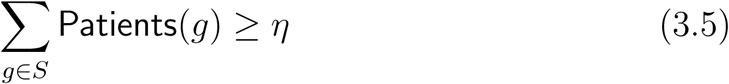

*where the counting g* ∈ *S is done without redundancy.*

The following definition is also valid for the persistent pathway complex *K*_*η*_.

#### Definition 4

*The space of pathways of a persistent pathway complex K*_*ε*_ *is the nerve generated by the facets of the complex, that is, the simplicial complex where* {*i*_0_, …, *i*_*ℓ*_} *is a simplex if and only if the facets indexed with i*_0_, …, *i*_*ℓ*_ *have a non empty intersection. We denote the space of pathways by* 𝒩_*ε*_.

The space of pathways is visualized through its 1-skeleton: the graph with pathways as vertices, and two pathways are connected if they intersect (see Figures 3 and 4).

**Figure 2:**
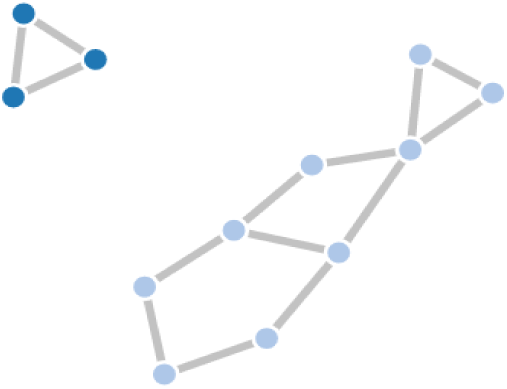
The graph has two connected components, which implies *β*_0_ = 2. It also has two cycles that are not triangles; thus, *β*_1_ = 2. Higher Betti numbers are zero.

**Figure 3:**
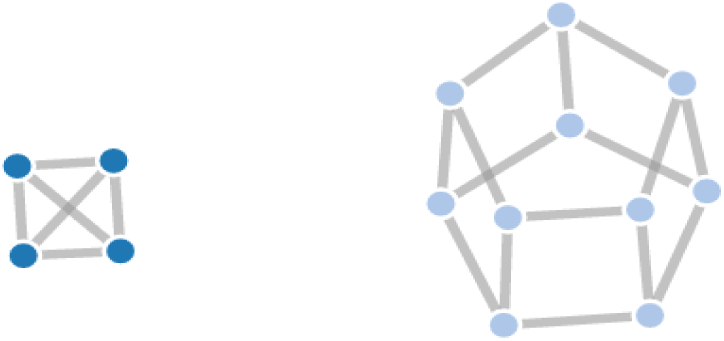
The graph has two connected components, giving *β*_0_ = 2. It also has 7 cycles that are not triangles, which yields *β*_1_ = 7. Higher Betti numbers are zero here as well. Now, if the different sides of the hexagonal prism (right component) are covered with triangles, then we get instead *β*_1_ = 0 and *β*_2_ = 1.

**Figure 4:**
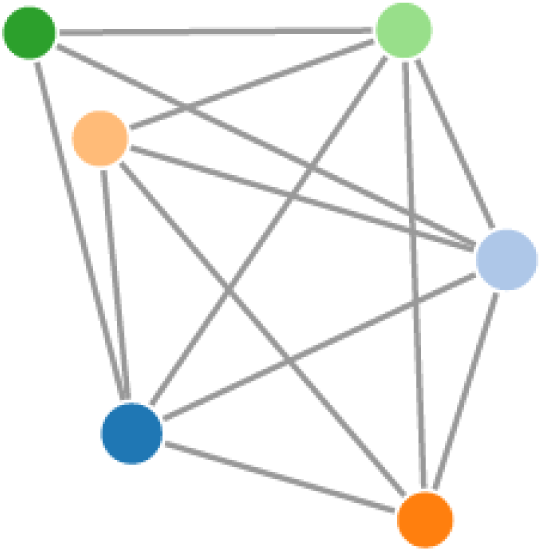
The 1-skeleton of the nerve 𝒩_*ε*_ for *ε* = 1 for AML data. In this case, the nerve is homotopy equivalent to the sphere. The different pathways represented by the nodes are given in the table above.

### 3.3 A primer on persistent homology

Recall that our plan is to compute the persistent homology of the independence complex *K*_*ε*_. We have explained that this is simply computing the homology of *K*_*ε*_ for a range of increasing values of the parameter *ε*. In this section, we explain this notion of homology (which we have introduced as measurements of the shape of the space *K*_*ε*_). For simplicity, we drop out the subscript *ε* from the complex *K*_*ε*_. It is also more convenient to introduce homology for clique complexes (for independence complexes, it suffices to replace, everywhere below, cliques with independent sets).

The homology of the simplicial complex *K* (now a clique complex of *G*) is a sequence of ℤ-vector spaces (i.e., vector spaces with integer coefficients):

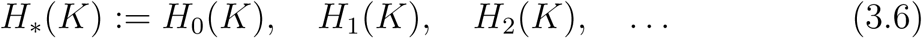

defined as follows:

- The zeroth vector space *H*_0_(*K*) is spanned by all connected components of *K*; thus, the dimension *β*_0_: = dim(*H*_0_(*K*)) gives the number of connected components of the space.
- The first homology space *H*_1_(*K*) is spanned by all closed chains of edges (cycles) in *G* which are not triangles – see Figure 2; in this case, the dimension *β*_1_: = dim(*H*_1_(*K*)) gives the number of “holes” in the space.
- Similarly, the second space *H*_2_(*K*) is spanned by all 2-dimensional enclosed three dimensional “voids” that are not tetrahedra (as in Figure 3 below).

Higher dimensional spaces are defined in a similar way (although less visual). Their dimensions count non trivial high dimensional voids. The dimensions *β*_*i*_: = dim(*H*_*i*_(*K*)) are called Betti numbers and provide the formal description of the concept of shape measurements.^4^

We move now to the notion of persistent homology and make it a bit more precise. For that, let us reintroduce the persistent parameter *ε* and let *K*_*ε*_ be again an independence complex. It is clear that if |Patients(*g*)∩Patients(*g*′)| ≥ *ε*_1_ and *ε*_1_ ≥ *ε*_2_ then the pair (*g, g*′), which is an edge in 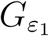, is also is an edge in 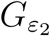. This means that 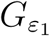 is a subgraph of 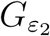; thus, we have 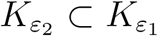 whenever *ε*_1_ ≥ *ε*_2_ (since an independent set for a given graph is also an independent set for any of its subgraphs). The mapping *ε* ↦ *K*_*ε*_ is functorial. It turns out that homology itself is functorial and all this functoriality is the mathematical reason that the following is correct: one can track the Betti numbers over a range of values *ε*_1_ ≥ *ε*_2_ ≥ *ε*_3_ ≥ … and consider the subrange where the Betti numbers are not changing (significantly). Pathways within this subrange are considered to have passed our test and declared robust computations.

## 4 Real mutation data

We have applied our approach to two mutation data (formulations and algorithms are available in [1]): Acute myeloid leukemia [14] and Glioblastoma multiforme [13]. For both data, we have computed the assignment *tumor* ↦ *pathways* through persistent pathway complexes (thus, declared robust output). The complete result is presented in long tables given in the Appendix. Interestingly, our calculation also shows that AML data is homotopy equivalent to a sphere while GBM data is homotopy equivalent to figure eight (genus-2 surface).

### 4.1 Acute myeloid leukemia data

The data has a cohort of 200 patients and 33 genes ([14]). We have chosen the coverage threshold *η* = 80 patient. We also neglected all genes that have fewer than 6 patients. These numbers are chosen based on the stability of barecodes for pairs (*ε, η*) ≥ (6, 80), while barecodes for pairs less than (6, 80) exhibit strong variations. This is also consistent with the fact that choosing genes with fewer than 5 or 6 patients is not common in such studies (genes with low numbers of patients are not considered robust enough, and are very prone to errors. This extra precaution is commonly used in the field). Now for the numbers of patients and coverage we have chosen, the Betti numbers are computed for various values of *ε* in the table below:

| $\varepsilon$ | $\vert \mathcal{N}_\varepsilon\vert $ | $density(\mathcal{N}_\varepsilon)$ | $\beta_i$ |
| --- | --- | --- | --- |
| 1 | 6 | 0.86 | $1, 0, 0, \dots$ |
| 2 | 84 | 0.97 | $1, 0, 0, \dots$ |
| 3 | 50 | 1 | $1, 0, 0, \dots$ |

Figure 4 below gives the 1-skeleton of the nerve 𝒩_*ε*_: = 𝒩(*K*_*ε*_) for *ε* = 1. The Betti numbers *β*_*i*_ are not changing; thus, *ε* = 1 is a reasonable choice. Recall that each node represents a pathway and two pathways are connected if they intersect (as sets of genes). We have used different colors to represent different pathways as described in the table below (no other meaning for the coloring).

| Color | Genes in the pathway |
| --- | --- |
| Blue | 'PML.RARA', 'MYH11.CBFB', 'RUNX1.RUNX1T1', 'TP53', 'NPM1', 'RUNX1' |
| Blue light | 'PML.RARA', 'MYH11.CBFB', 'RUNX1.RUNX1T1', 'TP53', 'NPM1', 'MLL.PTD' |
| Orange | 'PML.RARA', 'MYH11.CBFB', 'RUNX1.RUNX1T1', 'DNMT3A' |
| Orange light | 'Other Tyr kinases', 'MYH11.CBFB', 'MLL.PTD', 'NPM1' |
| Green | 'MLL-X fusions', 'TP53', 'FLT3' |
| Green light | 'Other Tyr kinases', 'MYH11.CBFB', 'DNMT3A', 'MLL-X fusions' |

### 4.2 Glioblastoma multiforme data

The second mutation data is taken from [13]. It has 84 patients and around 100 genes. Approximately, 70% of the genes have very low coverage so we removed them from the data; precisely, we have removed all genes with fewer than 10 patients. We have used the complex *K*_*η*_, with *ε* fixed to 7 because lower values don’t exhibit stable topologies; that is, barcodes are essentially one point long. This is a different choice of complex from the complex *K*_*ε*_ that we used with the previous AML data, which interestingly doesn’t yield noticeable stability–a possible explanation of this is the small size of the data (i.e., number of patients). Recall that the definitions of the two complexes are given on page 8 (definitions 2 and 3). In the table below, one can see that the topology stabilizes for the first three values of *η*. Any choice of pathways within this range is considered robust (proportionate to the small size of the data).

| $\eta$ | $\vert \mathcal{N}_\eta\vert $ | $density(\mathcal{N}_\eta)$ | $\beta_i$ |
| --- | --- | --- | --- |
| 66 | 15 | 0.73 | 1, 2, 0, … |
| 67 | 14 | 0.74 | 1, 2, 0, … |
| 68 | 12 | 0.72 | 1, 2, 0, … |
| 69 | 6 | 0.6 | 1, 0, 0, … |
| 70 | 50 | 1 | 1, 0, 0, … |

The following table provides the legend for the Figure 5 corresponding to the GBM data:

**Figure 5:**
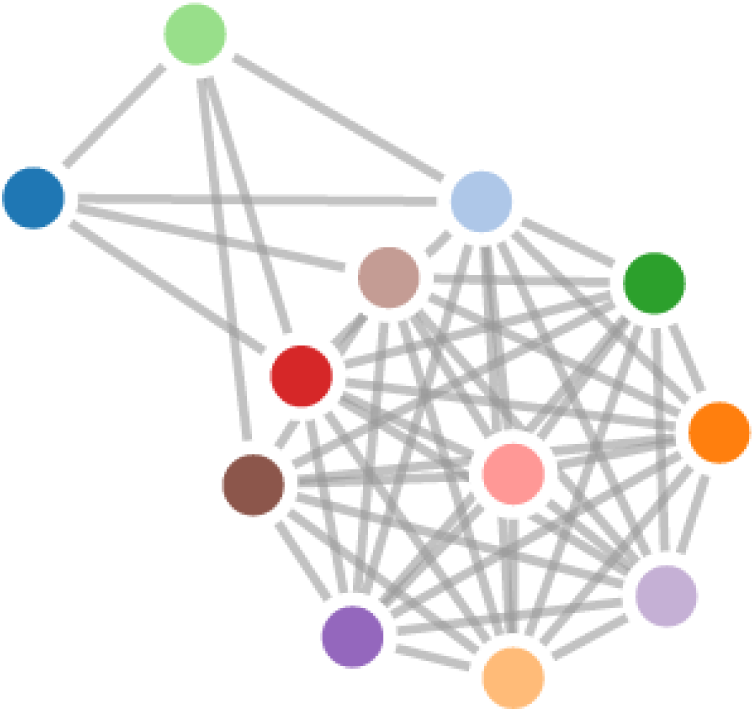
The nerve for the space of pathways for GBM data (see table below for the legend). The barcodes are stable in the first part of the table, which indicates that the nerve is homotopy equivalent to a genus-2 surface.

| Color | Genes in the pathway |
| --- | --- |
| blue | RB1, NF1, CYP27B1, CDKN2B |
| blue light | RB1, NF1, MDM2, AVIL-CTDSP2, CDKN2B |
| orange | TP53, MDM2, OS9, CDKN2A |
| orange light | TP53, MDM2, AVIL-CTDSP2, CDKN2A |
| green | TP53, MDM2, DTX3, CDKN2A |
| green light | RB1, NF1, CDK4, CDKN2B |
| brown | TP53, CDK4, CDKN2A |
| brown light | TP53, CYP27B1, CDKN2A |
| purple | TP53, MDM2, AVIL-CTDSP2, MTAP |
| purple light | TP53, MDM2, DTX3, MTAP |
| red | RB1, NF1, MDM2, OS9, CDKN2B |
| pink | TP53, MDM2, OS9, MTAP |

## 5 Conclusion

The main goal of this paper was to suggest a study of the *space of cancer pathways*, using the natural language of algebraic topology. We hope that the consideration of the pathways collectively, that is, as a topological space, helps to reveal novel relations between these pathways. Indeed, we have seen that the homology in the case of AML indicates that the mutation data has the shape of a sphere.^5^ However, in the case of GBM, the final set of pathways has the topology of a double torus (or, more technically, a genus-2 surface). This intriguing observation raises the question of whether these facts translate into a new biological understanding about cancer. Studying the space of pathways of other cancers will be illuminating as well, if they also show similar structures, and we can classify cancers by the topology of their mutated driver pathways. This is an example of the new type of hypotheses one can now formulate about the data. Eventually, our goal (recalling Poincare) is to help build a house by revealing patterns among the stones. Let us close with a quote from *The Emperor of All Maladies* (page 458):

> The third,^6^ and arguably most complex, new direction for cancer medicine is to integrate our understanding of aberrant genes and pathways to explain the *behavior* of cancer as a whole, thereby renewing the cycle of knowledge, discovery and therapeutic intervention.

## A Appendix

This appendix has three parts. In the first part, we review an efficient procedure for computing homology groups. The remaining two parts give the obtained list of pathways for GBM and AML data, respectively.

### A.1 Computing homology on quantum computers

We provide the following material (from [6]) for easy access. We review how the homology spaces *H*_*_(*X*) are computed. In principle, the formal definitions of homology that can be found in any algebraic topology textbook, are sufficient for computations. However, we point here to a more efficient approach based on Mayer-Vietoris blow-up complexes. We formulate finding optimal Mayer-Vietoris blow-up complexes as Quadratic Unconstrained Binary Optimizations (QUBO). QUBO-solvers (such as the D-Wave quantum annealer [8] or quantum-inspired classical solvers such as [2]) are now available for calculations.

Let 𝒞 = {*K*^*i*^}_*i*∈*I*_ be a cover of *K* by simplicial subcomplexes *K*^*i*^ ⊆ *K*; here again we have dropped the subscript *ε* from *K*_*ε*_ for simplicity. For *J* ⊆ *I*, we define 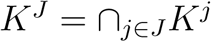.

#### Definition 5

*The* Mayer-Vietoris blow-up complex *of the simplicial complex K and cover* 𝒞, *is defined by:*

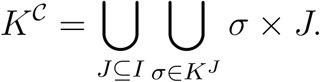

*A basis for the k*−*chains C*_*k*_(*K*^𝒞^) *is* {*σ* ⊗ *J* ∈ *K*^𝒞^| dim *σ* + card *J* = *n*}. *The boundary of a cell σ* ⊗ *J is given by: ∂*(*σ* ⊗ *J*) = *∂σ* ⊗ *J* + (−1)^d*im σ*^*σ* ⊗ *∂J.*

Simply put, the simplicial complex *K*^𝒞^ is the set of the “original” simplices in addition to the ones we get by blowing up common simplices. These are of the form *σ* ⊗ *J* in the definition above. In Figure 6, the yellow vertex *d* common to the two subcomplexes {*K*_1_, *K*_2_} is blown-up into an edge *d* ⊗ 12, and the edge *bc* is blown-up into the “triangle” *bc* ⊗ 01. In Figure 7, the vertex *a* common to three subcomplexes {*K*_0_, *K*_1_, *K*_2_} is blown-up into the triangle *a* ⊗ 012.

**Figure 6:**
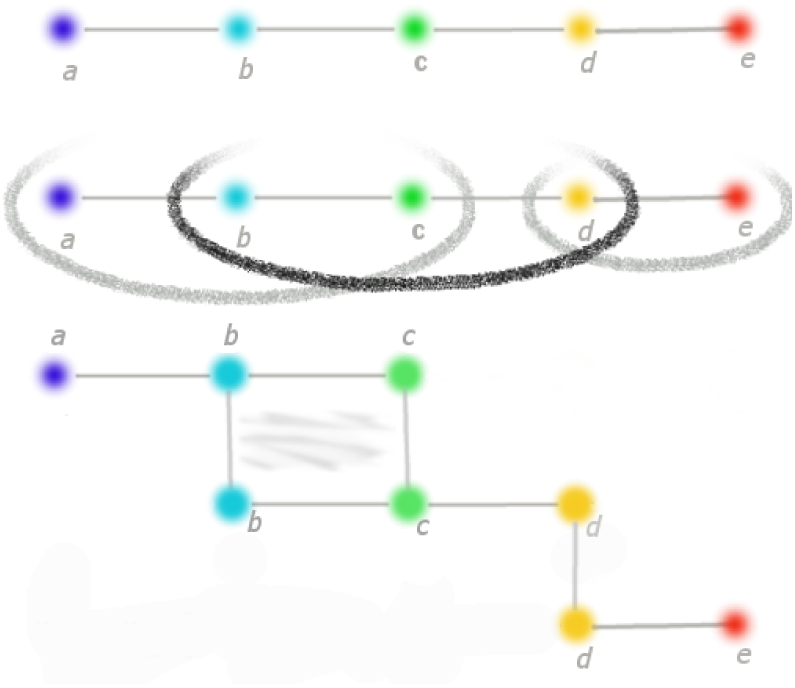
(Top) The simplicial complex *K* is the depicted graph. (Middle) *K* is covered with *K*_0_, *K*_1_, and *K*_2_. (Bottom) The blow-up complex of the cover depicted in the middle picture. After the blow-up, the edges *b*⊗01, *c*⊗01, *d* ⊗ 12, and the triangle *bc* ⊗ 01 appear.

**Figure 7:**
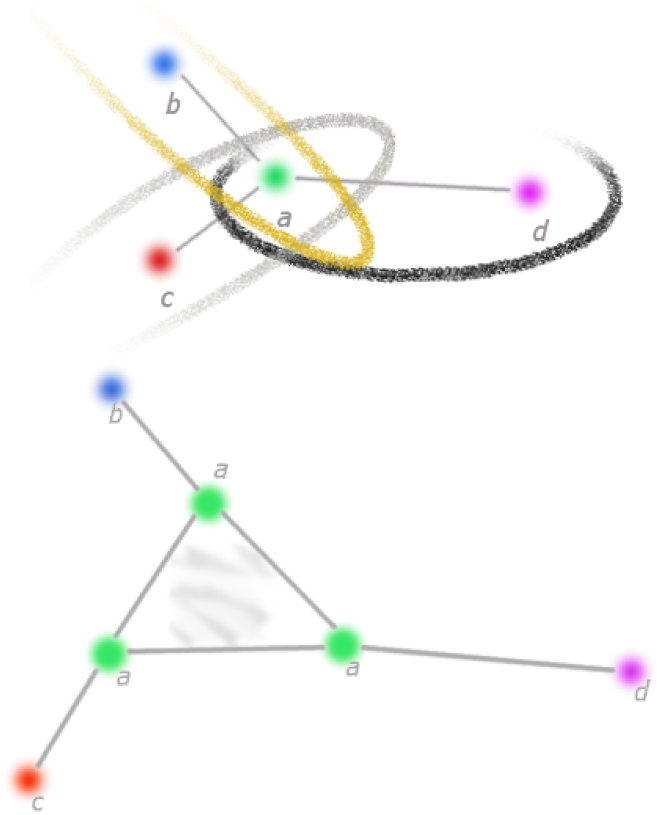
The triangle *a* × 012 appears after blowing-up the cover of the middle picture.

We will not prove it here, but the projection *K*^𝒞^ → *K* is a homotopy equivalence and induces an isomorphism *H*_*_(*K*^𝒞^) ≃ *H*_*_(*K*) [27]. The key point is that the boundary map of the simplicial complex *K*^𝒞^ (which replaces *K* by the homotopy equivalence) has a nice block form suitable for parallel rank computation. As an example, let us consider again the simplicial complex *K* depicted in Figure 6. First, *C*_0_(*K*^𝒞^), the space of vertices of the blow-up complex *K*^𝒞^, is spanned by the vertices

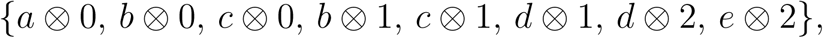

that is, all vertices of *K* taking into account the partition they belong to. The space of edges *C*_1_(*K*^𝒞^) is spanned by

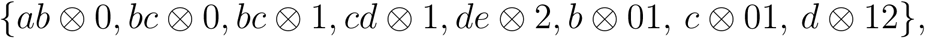

which is the set of the “original” edges (edges of the form *σ* ⊗ *j* with *j* ∈ *J* = {0, 1, 2} and *σ* is an edge in *K*) and the new ones resulting from blowups, that is, those of the form *v* ⊗ *ij* where *v* is a vertex in *K*^*j*^ ∩ *K*^*j*^ (if the intersection is empty, the value of boundary map is just 0). The matrix of the boundary map *∂*_0_ with respect to the given ordering is then:

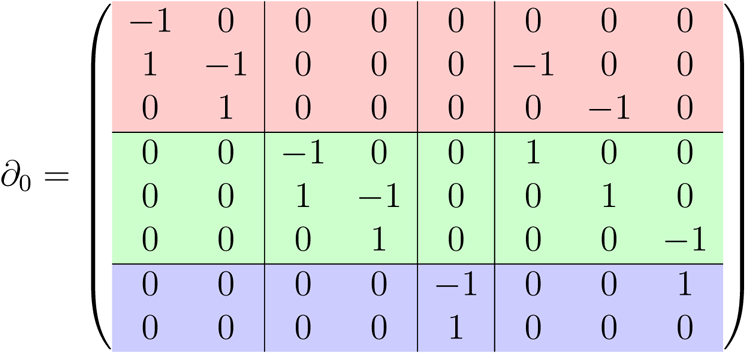

Clearly, one can now row-reduce each coloured block independently. There might be *remainders*, that is, zero rows except for the intersection part. We collect all such rows in one extra matrix and row-reduce it at the end and aggregate. For the second boundary matrix we need to determine *C*_2_(*K*^𝒞^). The 2-simplices are of three forms. First, the original ones (those of the form *σ* ⊗ *j* with *σ* ∈ *C*_2_(*K*); in this example there is none) then those of the form *σ* ×{*i, j*}, with *σ* being in *K*^*i*^ ∩ *K*^*j*^. And finally, those of the form *v* ⊗{*i, j, k*}, with *v* ∈ *K*^*i*^ ∩ *K*^*j*^ ∩ *K*^*k*^ (there is none in this example, but Figure 5 has one). We get *C*_2_(*K*^𝒞^) = ⟨*bc* ⊗ 01⟩; thus, there is no need for parallel computation.

Finally, for efficient computations, it is necessary that the homologies of the smaller blocks are easy to compute. This is where the QUBO-solver comes in. It is used to obtain a “good” cover 𝒞 by computing a clique cover of the original graph *G* (plus a completion step). It is easy to see that such coverings do, indeed, come with trivial block homologies [6].

### A.2 GBM pathways list

The table below reports the final pathways in Figure 5. Details on the data, the parameters, and how the pathways are computed using persistent homology, can be found in Subsection 4.2 on page 13.

| Pathway | coverage $\eta$ |
| --- | --- |
| RB1 , NF1 , CYP27B1 , CDKN2B | 70 |
| RB1 , NF1 , MDM2 , AVIL-CTDSP2 , CDKN2B | 69 |
| TP53 , MDM2 , OS9 , CDKN2A | 69 |
| TP53 , MDM2 , AVIL-CTDSP2 , CDKN2A | 69 |
| TP53 , MDM2 , DTX3 , CDKN2A | 69 |
| RB1 , NF1 , CDK4 , CDKN2B | 69 |
| RB1 , NF1 , MDM2 , OS9 , CDKN2B | 68 |
| TP53 , MDM2 , OS9 , MTAP | 68 |
| TP53 , MDM2 , AVIL-CTDSP2 , MTAP | 68 |
| TP53 , MDM2 , DTX3 , MTAP | 68 |
| TP53 , CDK4 , CDKN2A | 68 |
| TP53 , CYP27B1 , CDKN2A | 68 |
| TP53 , CDK4 , MTAP | 67 |
| TP53 , CYP27B1 , MTAP | 67 |
| RB1 , NF1 , CYP27B1 , CDKN2A | 66 |
| RB1 , NF1 , MDM2 , OS9 , CDKN2A | 65 |
| RB1 , NF1 , MDM2 , AVIL-CTDSP2 , CDKN2A | 65 |
| RB1 , NF1 , MDM2 , DTX3 , CDKN2B | 65 |
| RB1 , NF1 , CDK4 , CDKN2A | 65 |
| RB1 , NF1 , CYP27B1 , MTAP | 65 |
| RB1 , NF1 , MDM2 , OS9 , MTAP | 64 |
| RB1 , NF1 , MDM2 , AVIL-CTDSP2 , MTAP | 64 |
| RB1 , NF1 , CDK4 , MTAP | 64 |
| RB1 , NF1 , MDM2 , DTX3 , CDKN2A | 62 |
| RB1 , NF1 , MDM2 , DTX3 , MTAP | 61 |
| TP53 , MDM2 , AVIL-CTDSP2 , SEC61G | 60 |
| RB1 , NF1 , EGFR , OS9 | 60 |
| TP53 , MDM2 , OS9 , SEC61G | 59 |
| TP53 , MDM2 , DTX3 , SEC61G | 59 |
| PTEN , IFNA21 , MDM2 , AVIL-CTDSP2 | 58 |
| TP53 , MDM2 , AVIL-CTDSP2 , IFNA21 | 58 |
| RB1 , NF1 , EGFR , DTX3 | 58 |
| PTEN , IFNA21 , CYP27B1 | 58 |
| PTEN , IFNA21 , MDM2 , OS9 | 57 |
| TP53 , MDM2 , OS9 , IFNA21 | 57 |
| TP53 , MDM2 , DTX3 , IFNA21 | 57 |
| PTEN , IFNA21 , CDK4 | 57 |
| RB1 , NF1 , MDM2 , AVIL-CTDSP2 , SEC61G | 55 |
| PTEN , IFNA21 , MDM2 , DTX3 | 55 |
| TP53 , MDM2 , AVIL-CTDSP2 , ELAVL2 | 55 |
| RB1 , NF1 , MDM2 , OS9 , SEC61G | 54 |
| TP53 , MDM2 , OS9 , ELAVL2 | 54 |
| TP53 , MDM2 , DTX3 , ELAVL2 | 54 |
| RB1 , NF1 , MDM2 , AVIL-CTDSP2 , IFNA21 | 53 |
| TP53 , CDK4 , IFNA21 | 53 |
| TP53 , CDK4 , ELAVL2 | 53 |
| TP53 , CYP27B1 , IFNA21 | 53 |
| TP53 , CYP27B1 , ELAVL2 | 53 |
| RB1 , NF1 , MDM2 , OS9 , IFNA21 | 52 |
| RB1 , NF1 , MDM2 , DTX3 , SEC61G | 52 |
| RB1 , NF1 , MDM2 , AVIL-CTDSP2 , ELAVL2 | 51 |
| RB1 , NF1 , CYP27B1 , IFNA21 | 51 |
| RB1 , NF1 , CYP27B1 , ELAVL2 | 51 |
| RB1 , NF1 , MDM2 , OS9 , ELAVL2 | 50 |
| RB1 , NF1 , CDK4 , IFNA21 | 50 |
| RB1 , NF1 , CDK4 , ELAVL2 | 50 |
| RB1 , NF1 , MDM2 , DTX3 , IFNA21 | 49 |
| RB1 , NF1 , MDM2 , DTX3 , ELAVL2 | 47 |

### A.3 AML pathways list

The following table gives the list of all 65 pathways of 𝒩_1_ (Figure 4) and their coverage. Details about the data and how the pathways are obtained using persistent homology can be found in Section 3.1. All pathways listed below passed our robustness test.

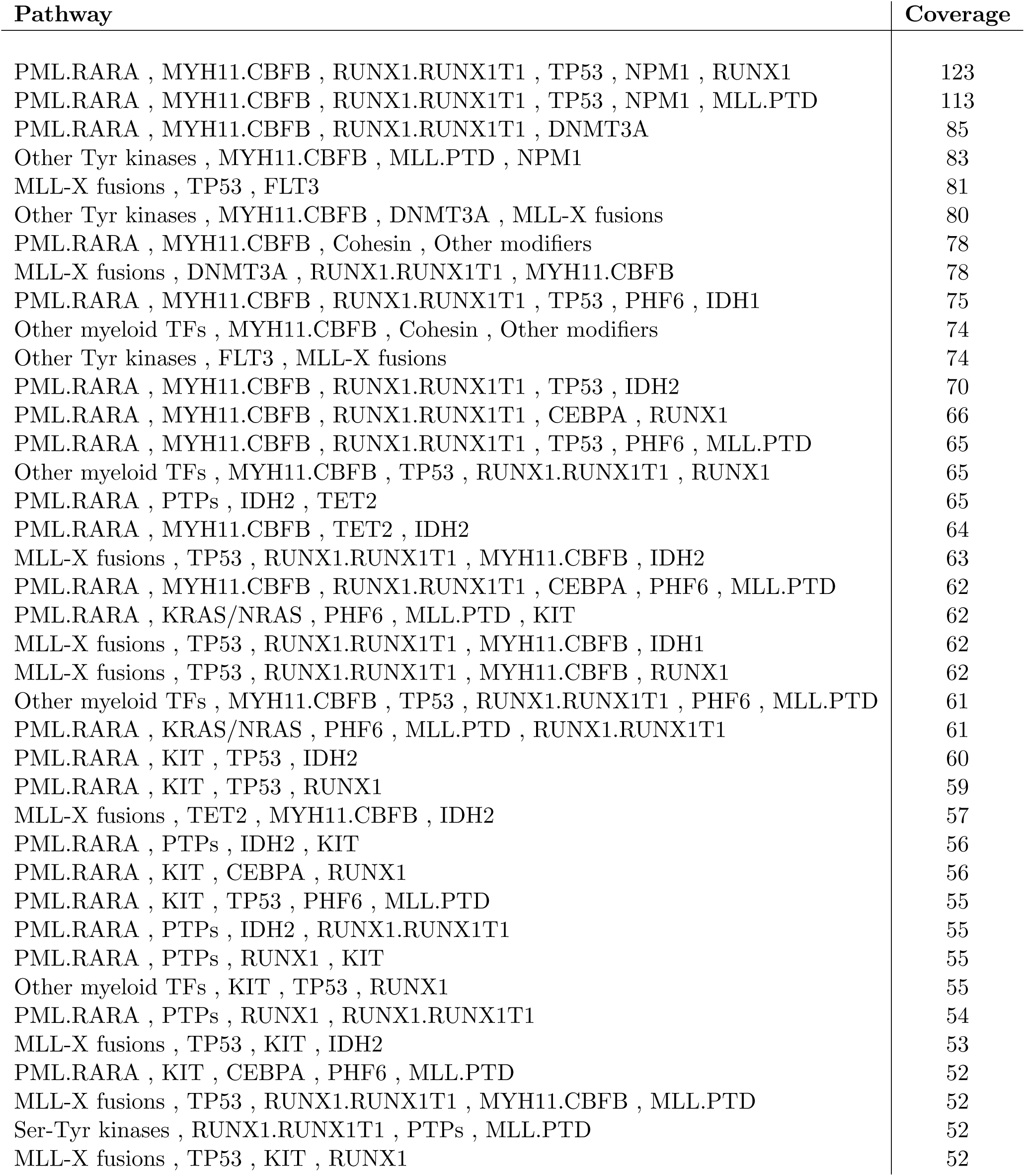

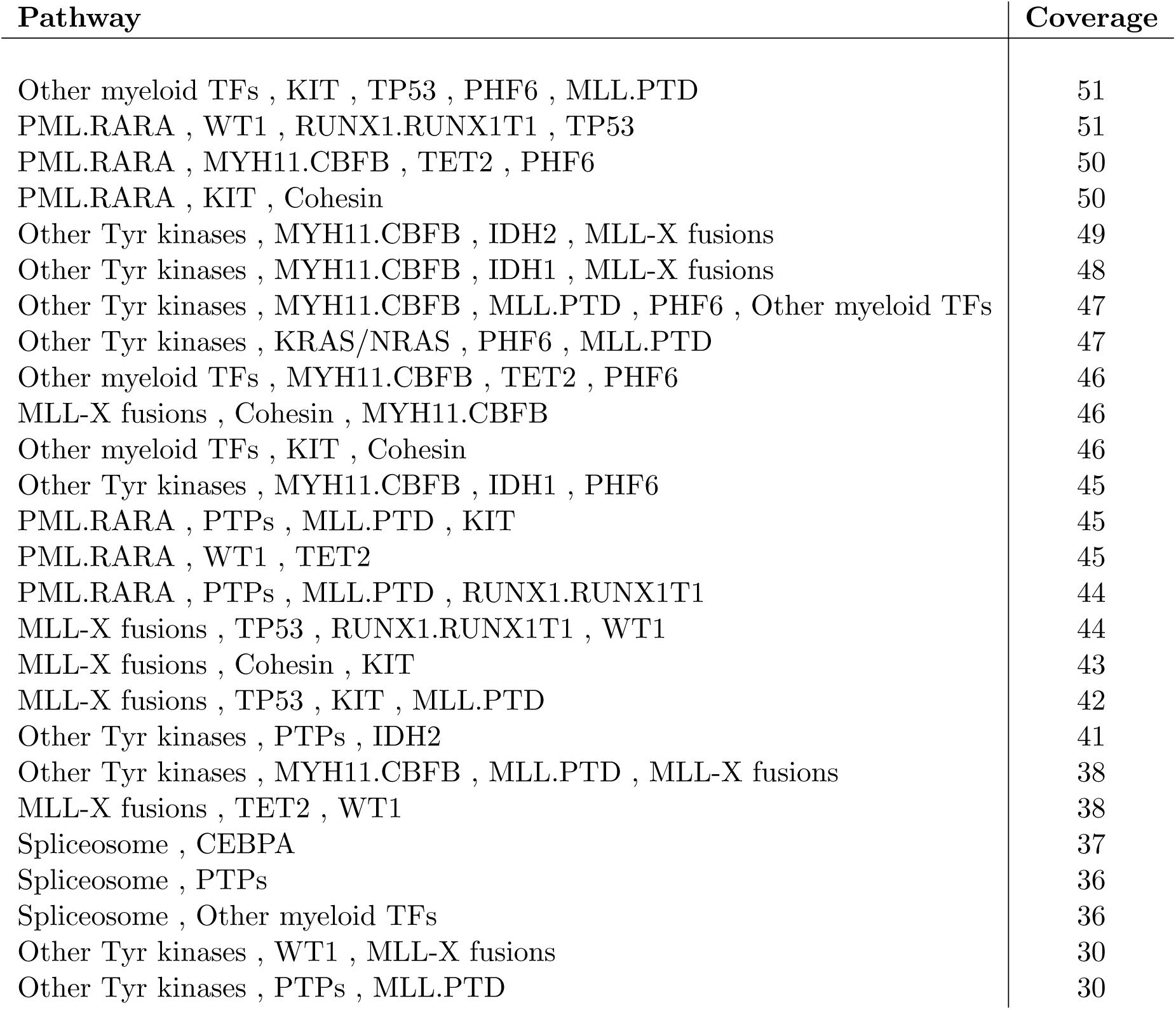

The following long table gives the assignment *tumor* ↦ *list of pathways* for the AML data. For each tumor we assign a robust (in our topological sense) set of pathways. More specifically, for a tumor (sample ID) i, the second column gives the list of all pathways that have passed our robustness test.

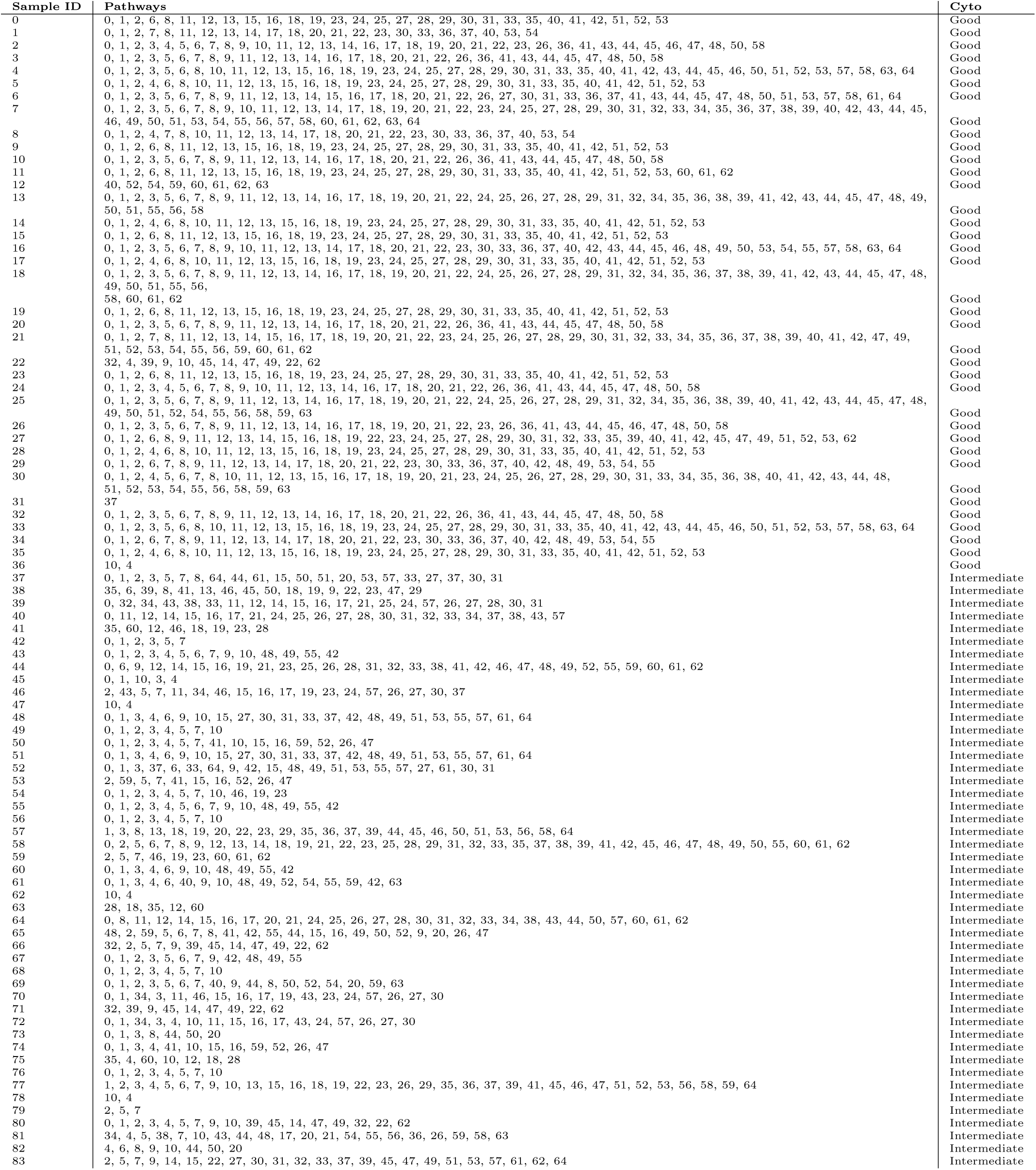
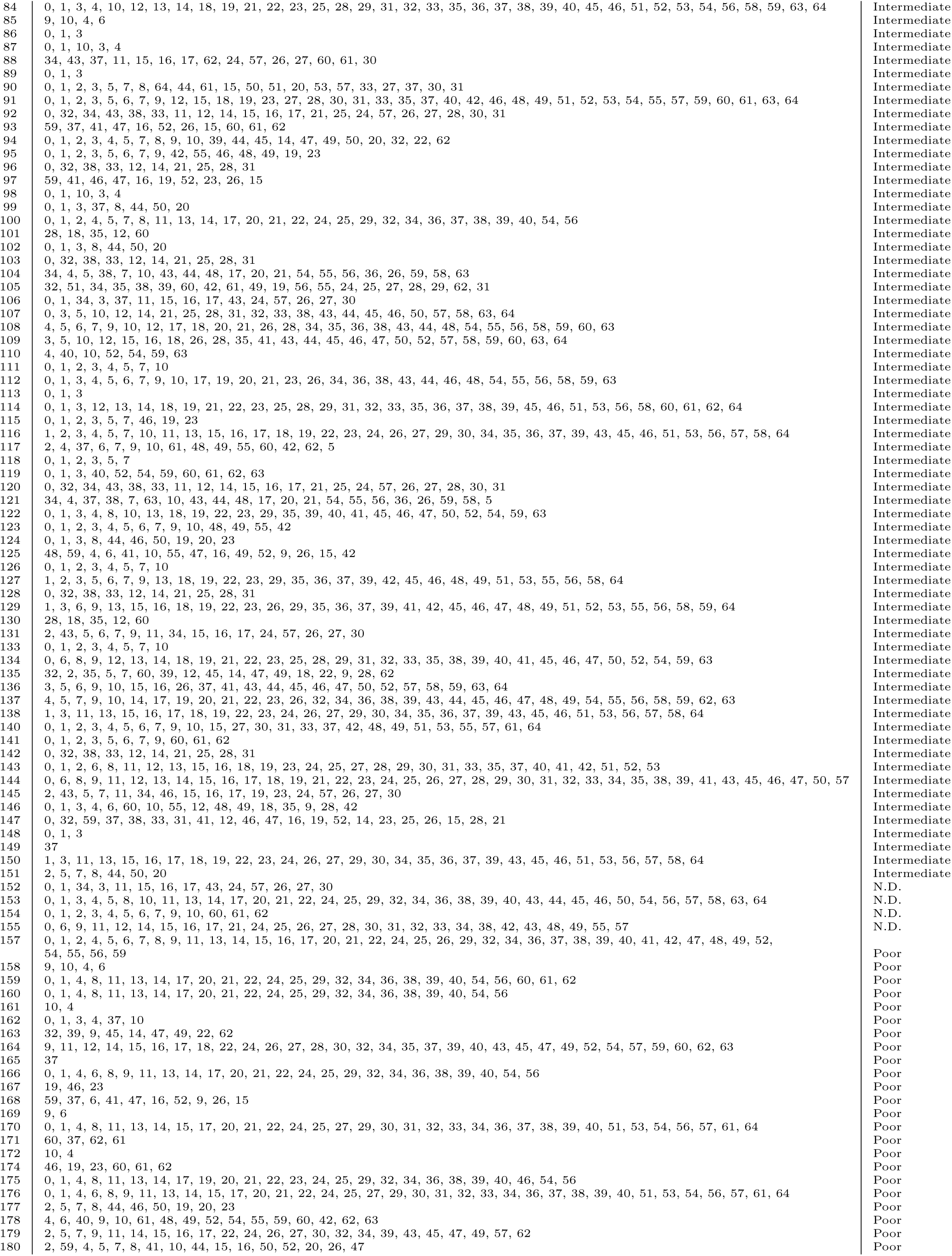
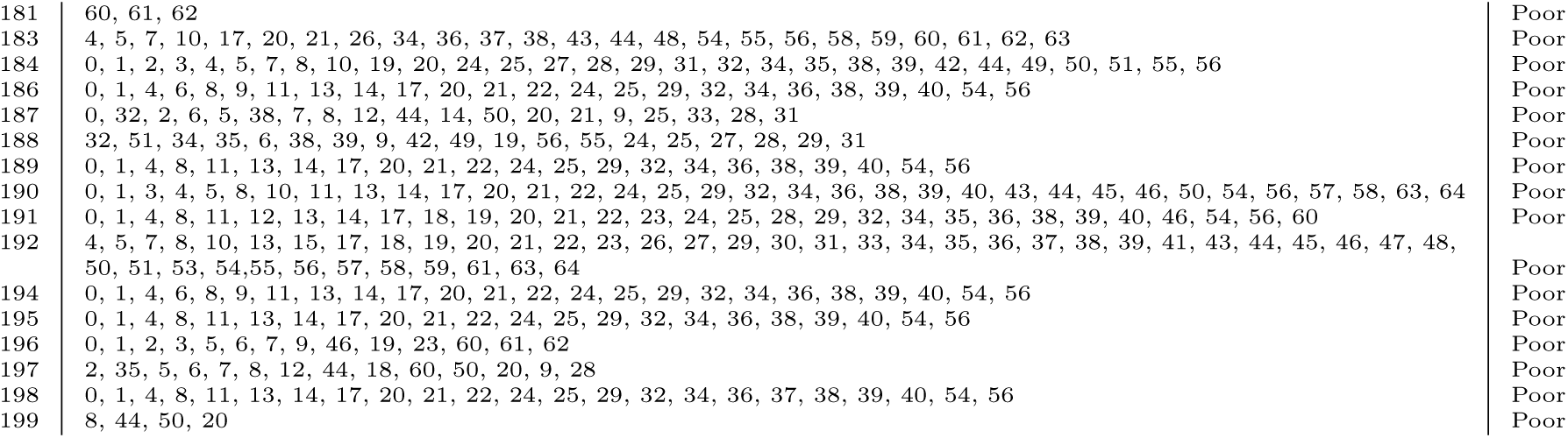

## Footnotes

1 Current advances include the introduction of a single anti-cancer agents, which simply bind with growth factor receptors stopping abnormal cell proliferations. For instance, in the context of breast cancer, Herceptin antibody stops cells abnormal proliferation signals by binding with the excess of growth factor receptors on the cell surface caused by point mutations in the gene Her2. Gleevec is a second example. It is used to treat chronic myeloid leukemia, which is a type of blood cell tumor due to an inappropriate gene fusion product of a translocation in chromosomes 9 and 22. The resulting fusion gene BCR-ABL has an increased kinase activity, resulting in an increase in proliferation signals. Similar to Herceptin, Gleevec blocks the growth signals that the abnormal fusion gene generates and thus prevents the cell proliferation. More sophisticated approaches include immunotherapy that activates the immune system against cancer cells as well as the use of combinations of therapeutic agents to attack multiple pathways fundamental in cancer development, preventing resistance from occurring.

2 In the language of *Homotopy Type Theory (HoTT)* [19], this procedure translates into: *tumor* (∞, 1) *category*. This ↦ (∞,1)−category is an (∞,1)−topos (a logical structure) in the sense of Lurie [12], so it obeys the *axioms* of HoTT.

3 Commercial applications of this approach in various different areas of practice are tackled through Ayasdi.

4 In textbooks, one typically starts with a continuous space (for instance, a doughnut shaped surface) and then triangulizes it, yielding the simplicial complex *K*. The homology of the continuous space is the homology of its discretization *K* that we introduced pictorially above. Appendix A, on page 19, explains how to compute homology using quantum computers or QUBO-solvers such as [2].

5 Using a different visual representation, Vogelstein found AML to be very different from other cancers. Indeed, that has allowed many scientists to speculate that such genetically simple tumors are more susceptible to drugs, and thus intrinsically more curable.

6 The first is targeted therapy on the mutated pathways, as we mentioned in the Introduction. The second is cancer prevention through identifying preventable carcinogens.

